# Nuclear proteome analysis reveals spatial and temporal compartmentalization of metabolic enzymes during *Trypanosoma cruzi* cell cycle

**DOI:** 10.1101/2025.06.30.662266

**Authors:** A.P.J. Menezes, C. Gachet-Castro, S. McGill, R. Burchmore, A.M Silber, M.C Elias, J.P.C. Cunha

## Abstract

*Trypanosoma cruzi*, the etiological agent of Chagas Disease (CD), represents a significant public health concern and serves as a valuable model for investigating the cell cycle in early-diverging eukaryotes. Its unique cellular features—including the absence of chromosome condensation and the coordination of nuclear division with specialized organelles—provide insights into non-canonical regulatory mechanisms, and underscore the pivotal role of the nucleus in maintaining genomic integrity and regulating fundamental processes such as DNA replication and gene expression. To investigate nuclear dynamics throughout the cell cycle, we synchronized *T. cruzi* epimastigotes at G1/S, S, and G2/M using hydroxyurea (HU), and isolated nuclear proteins were quantitative evaluated by LC-MS/MS. We identified 2,937 nuclear-enriched proteins, revealing distinct, phase-specific expression patterns. The G1/S phase was marked by increased levels of metabolic enzymes including those related to energy and nucleotide/nucleoside metabolisms. The S phase showed elevated abundance of canonical and variant histones, consistent with chromatin remodeling and DNA replication. The G2/M phase was enriched in proteins involved in protein synthesis and microtubule dynamics, essential for mitosis. Notably, ∼6% of the nuclear proteome consisted of metabolic enzymes, a finding further supported by other public proteomics datasets and in vitro nuclear activity assays. These results suggest that metabolic enzymes and/or their metabolites may modulate nuclear processes in *T. cruzi*, potentially influencing the epigenome and gene expression regulation, as proposed in other eukaryotic models. Altogether, our findings reveal dynamic remodeling of the nuclear proteome during the *T. cruzi* cell cycle and point to a previously underappreciated role for metabolic enzymes likely regulating nuclear functions in a cell cycle-dependent manner.

## 1. Introduction

The *Trypanosoma cruzi*, a protozoan of Trypanosomatidae order, is the etiological agent of Chagas Disease (CD), a neglected tropical disease with significant morbidity and mortality (1,2). According to the World Health Organization, it is estimated that 8 million people are infected with *T. cruzi* (2), primarily in Latin America (3). In recent decades, there has been an increase in cases in non-endemic countries due to migratory flows in a globalized context, making CD an emerging health problem (4).

In addition to its medical relevance, *T. cruzi* serves as a valuable model for research due to its classification as an early diverging eukaryote (5). It exhibits distinctive characteristics, including polycistronic transcription (6), RNA editing (7), and the presence of specialized organelles such as the glycosome (8). The cell division cycle of *T. cruzi* exhibits distinctive features, including the coordination of nuclear division with single-copy organelles such as the basal body, the flagellum, and the mitochondrion, which contains the kinetoplast — a network of circular DNA molecules (9). Cell cycle progression is tightly coupled to the emergence and elongation of a new flagellum and to the division of the nucleus (N) and kinetoplast (K). These coordinated events, which are distinguishable by microscopy, enable accurate staging of the cell cycle by microscopy: cells with 1N1K1F are in G1 or S phase; 1N1K2F corresponds to G2; and 2N1(2)K2F indicates mitosis or cytokinesis (10). DNA replication also displays differences relative to other eukaryotes. DNA replication occurs mainly at the nuclear periphery (11) and the pre-replication complex lacks some canonical proteins (12,13). Additionally, trypanosomes display a closed mitosis and lacks chromosome condensation (14). Consequently, in these organisms’ metaphase it is not possible to visualize the alignment of chromosomes in the equatorial cell region, as in other eukaryotes (15). Thus, the molecular mechanism governing cell cycle must be divergent from those described in other organisms. Therefore, comprehending *T. cruzi* cell cycle biology is crucial not only for elucidating ancestral cellular mechanisms that contribute to evolutionary processes, but also from a biomedical perspective. As a pathogenic organism, identifying critical regulators of its cell cycle may uncover novel therapeutic targets to control parasite replication and, consequently, disease. This is especially relevant considering that trypanosomatids are evolutionarily divergent eukaryotes, and the regulatory mechanisms uncovered may differ significantly from those found in the human host—offering opportunities for selective therapeutic intervention.

The cell cycle of *T. cruzi* has already been investigated by proteomics and transcriptomics assays. Transcriptomics cell cycle evaluation showed that the most regulated cell cycle transition was G2/M to G1, in which cellular functions related to the metabolism of carbohydrates and energy production were overrepresented (16). Whole proteomics assays of parasites in G1/S and S showed an up-regulation of proteins involved in oxidative phosphorylation while pathways related to protein synthesis decreased (17). We and others have been investigating the role of nuclear proteins as a central hub for regulating key cellular processes such as replication and transcription. Previously, we identified ∼ 2,200 chromatin-associated proteins in *T. cruzi*, while others reported 900 nuclear proteins, with a significant proportion (31%) of proteins with unknown function (18). Considering that, nuclear proteome analysis provides insights into protein-level regulation, which is particularly relevant in trypanosomatids, where gene expression is predominantly controlled post-transcriptionally (6), we aimed to characterize the nuclear subproteome across the cell cycle. Our study identified several hundred nuclear proteins, including both canonical and non-canonical components, many of which display dynamic changes across the cell cycle.

## 2. Material and Methods

### 2.1 Cell culture and cell cycle synchronization

*T. cruzi* epimastigotes (Cl Brener strain) were grown in liver infusion tryptose (LIT) medium supplemented with 10% heat-inactivated fetal bovine serum at 28 °C. Different cell cycle stages were obtained after synchronization as previously performed (11,19). Briefly, parasites in a log-phase were treated with 20 mM of Hydroxyurea (HU) for 24 h. HU was then removed from the medium by centrifugation (1200 x *g*, 10 min), the cells were washed three times in phosphate-buffered saline (PBS) and resuspended in fresh LIT medium. To obtain G1/S, S and G2/M enriched parasite populations, samples were collected at 0, 6 and 12 h post-HU release respectively. The samples were kept at −80 °C until the cell cycle synchronization confirmation. A sample of parasites harvested before the incubation with HU was used as a control of asynchronous growth. For each time-point, an aliquot of 10^7^ parasites/mL was washed twice in cold-PBS and fixed overnight at 4°C with 70% Ethanol-PBS. Propidium iodide (PI) staining was carried out by incubation of the fixed parasites for 30 min at 37°C in PBS containing 20 μg/mL PI and 20 μg/mL RNAse (Invitrogen) for DNA-specific staining. Three technical replicates per biological sample were analyzed for DNA-content in a Flow cytometer (Attune NXT - Thermo). The proportion of G1/S, S and G2/M cells in the samples was determined by DNA content analyzed in FlowJo software (BD).

### 2.2 Nuclei Purification

Initially, freshly parasites (1x10^9^) were rinsed and resuspended in 1 ml of a solution comprising 10 mM potassium glutamate, 250 mM sucrose, 2.5 mM CaCl_2_, and 1 mM phenyl-methylsulfonyl fluoride, designated as the suspension buffer. Subsequently, lysis was induced by the addition of 0.1 Triton X100. The resulting lysate underwent centrifugation, followed by washing and resuspension in 1 ml of suspension buffer. The lysate was then carefully layered atop a 50% Percoll solution (Sigma) in a suspension buffer (4 ml) and subjected to centrifugation at 63,000 × *g* at 4 °C for 2 hours, as previously described (20). The nuclei, located at the bottom of the self-forming Percoll gradient were isolated, washed thoroughly with a suspension buffer, and prepared for further analysis.

### 2.3 TMT labeling and quantitative proteomics analysis

Nuclei protein extracts were prepared with FASP™ Protein Digestion Kit (Expedeon-Abcam-USA) and digested with trypsin (Promega) by following the manufacturer’s recommendations. After the digestion of proteins into peptides, the samples were labeled with TMT® (Thermo-TMT 16plex Mass Tag Labeling kit cat. 90110). Briefly, 50 μg of peptides were resuspended to 40 μl of 100 mM Tetraethylammonium bicarbonate (TEAB) and 10 μl of 100 % acetonitrile. Ten microliters of the TMT label reagent were added to each sample, and the reaction was allowed to proceed for 1 hour at room temperature. For reaction quenching, we added 8 μl of 0.5 % hydroxylamine per sample and incubated for 15 minutes. Equal amounts of each sample (126, 127N, 127C, 128N, 128C, 129N, 129C, 130N,130C, 131N, 131C, 132N) were combined and dried in a speed vac. Peptides were separated and analyzed by an UltiMate 3000 RSLC nano system coupled to an Orbitrap Fusion Tribrid mass spectrometer (both from Thermo Scientific). Peptides were firstly loaded and concentrated on to an Acclaim PepMap300 trap column (300 µm x 5 mm) packed with C18 (5 µm, 300 Å) (Thermo) in loading buffer of 1 % acetonitrile with 0.1 % formic acid then separated in an EASY-Spray column (75 µm x 50 cm) packed with C18 (2 µm, 100 Å, Thermo Scientific) at a flow rate of 300 nL/min. Mobile phase A (0.1% formic acid in water) and mobile phase B (0.08 % formic acid in 80% acetonitrile / 20% H_2_O) were used to establish a solvent gradient of 4 % B for 1.5 min, 4 to 60 % for 178.5 min, 60 to 99 % for 15 min, and held at 99 % for 5 min. A further 10 minutes at initial conditions for column re-equilibration was used before the next injection. The Orbitrap Fusion acquired data continuously high-resolution precursor scan at 120 000 resolving power (over a mass range of m/z 400 – 1600) followed by top speed (2.5s) CID fragmentation (35 %) and detection of the top precursor ions from the MS scan in the linear ion trap using turbo scan speed. Triple stage mass spectrometry (MS3) higher-energy collisional dissociation (HCD; 55%) was performed on the top ions from the MS2 CID scan, with up to 10 synchronous precursor selection (SPS) scans isolated with the precursor ion and any TMT loss ions excluded from the selection. Orbitrap detection of the TMT quantitation label from the MS3 fragmentation was acquired at a resolution of 50000 over a mass range of 100-500 m/z.

### 2.4 TMT data processing

Thermo Scientific Proteome Discoverer (version 2.5) with SEQUEST® search engines were used for peptide/protein identification. The first step of the data processing was to check the TMT labeling efficiency (greater than 90%, there is less than a 3-fold difference in an individual channel). Reporter ion intensities in each TMT channel were aggregated to the peptide level. Mean values were then calculated for each identified peptide across all TMT channels, and the values for each channel were then divided by this mean. The resulting ratios were log_2_-transformed and used to make distribution plots for each TMT channel within a plex. Normalization was applied based on total peptide amount, with factors ranging from 1.0 to 2.46 for different samples (Supplementary Fig2B). These values were used to ensure consistent protein quantification across the TMT channels. The protein searches were performed against a *T. cruzi* database (*Trypanosoma cruzi* CL Brener Esmeraldo-like V.54, available on TriTrypDB). Proteins were grouped by applying the maximum parsimony rule (i.e., the protein groups in the final report represent the shortest possible list needed to explain all confidently observed peptides). The quantitation module within Proteome Discoverer™ software was used to assess the ratios for individually tagged samples. The height of reporter ions detected with mass tolerance ±10 ppm was adjusted, considering the isotopic correction factors provided by the TMT kit manufacturer. The searches were performed with the following parameters: MS/MS accuracy 0.6 Da for CID and 0.1 Da for HCD, trypsin digestion with two missed cleavages allowed, fixed carbamido-methyl modification of cysteine, and variable modification of oxidized methionine and phosphorylation (S, T, Y). TMT Pro was set as a variable modification of the N-terminus. The number of proteins and protein groups and numbers of peptides were estimated using Proteome Discoverer, with a false discovery rate (FDR) of 5%, and peptide rank 1 applied as a cutoff limit. Proteins showing fold changes ≥ 1.5 or ≤ −1.5 and and p-value < 0.05 for each set of comparisons were considered and displayed on volcano plots.

### 2.5 Gene Ontology (GO) and metabolic pathways enrichment and other *T. cruzi* proteomes

We conducted the GO enrichment analysis through the TriTrypDB database (https://tritrypdb.org/tritrypdb/app/) to identify significantly enriched GO terms among the selected proteins. We searched for the total proteins found in each cell cycle phase by cellular component category. Next, we examined the differentially expressed proteins (fold changes ≥ 1.5 or ≤ −1.5 and p-value < 0.05) in the ratios of G1/S vs. S and G2/M vs. G1/S to identify significantly enriched GO terms in biological processes and molecular functions. We set the threshold value for GO terms with a p-value cutoff of 0.05.

*T. cruzi* metabolic enzymes were classified as those belonged to the following GO terms available at TriTrypDB: “cellular amino acid metabolic process” (GO: 0006520), “carbohydrate metabolic process” (GO: 0005975), “pyruvate metabolic process” (GO: 0006090), “ATP metabolic process” (GO: 00046034), “nucleotide metabolic process” (GO: 0009116), “nucleoside metabolic process” (GO: 0009117), “lipid metabolic process” (GO: 0006629), “generation of precursor metabolites and energy” (GO: 0006091) terms. Metabolic pathway enrichment was conducted by KEEG tool though the TriTrypDB DB considering a p-value cutoff of 0.05 (Bonferroni). The list of proteins detected at the nuclear (18), chromatin (21) and cell cycle (17) proteomics was obtained from supplemental material and compared to our dataset.

### 2.6. Preparation of epimastigotes extracts for enzymatic assays

Enzymatic activity assays were performed with the samples obtained from whole extracts, nuclear and cytoplasmic extracts of epimastigotes forms. Parasites in the exponentially growing phase in LIT medium were collected and washed twice with chilled 1X PBS. Cell lysis was performed according to (22). Briefly, parasites in hypotonic buffer (10 mm HEPES, pH 7.9; 1.5 mm MgCl_2_; 10 mm KCl; 0.5 mm DTT; and protease inhibitors) with 0.1% NP40, and the cytoplasm was separated from the nucleus after the addition of 1 M sucrose and centrifugation at 500 x *g* for 5 minutes at 4°C. The supernatant containing the cytoplasm was collected and kept at 4°C, while the pellet was washed with a 0.35 M sucrose solution. After centrifugation at 1100 x g for 5 minutes at 4°C, the supernatant was discarded, and the nucleus was solubilized in a buffer containing 0.35 M sucrose, 0.5 mM DTT; 10 mM HEPES pH 7.9; 33 mM MgCl_2_; 10 mM KCl; and a cocktail of protease inhibitors (Sigma). The total extract was obtained by lysing the parasites with buffer containing 20 mM Tris-HCl, 0.25 M sucrose, 1 mM EDTA, 0.1% Triton-X 100 (v/v), 1 mM PMSF, and a cocktail of protease inhibitors (Sigma) followed by sonication for five cycles (5 seconds at 20% power and 30 seconds pause on ice). After this process, the extracts were centrifuged at 13,000 x *g* for 30 minutes at 4°C, and the supernatants were quantified using the Bradford method (23). Following the activity assays, total, cytoplasmic, and nuclear extracts were analyzed via Western Blotting (WB) to confirm the nucleus and cytoplasm separation, using anti-eIF5a antibodies (cytoplasmic protein) and anti-histone H3 antibodies (nuclear protein).

### 2.7. Enzymatic assays

Hexokinase (HK) activity assay: HK was assessed indirectly through a coupled enzymatic reaction involving glucose-6-phosphate dehydrogenase (G6PD, Sigma). In this assay, HK catalyzes the conversion of glucose and ATP into glucose-6-phosphate (G6P), which is subsequently oxidized by G6PD, generating NADPH. The formation of NADPH is monitored by measuring absorbance at 340 nm. Thus, in the presence of glucose and ATP, an increase in absorbance over time indicates HK activity, with a linear rise in NADPH production reflecting the enzymatic reaction rate. In this assay, the reaction contained: 50 mM Tris-HCl pH 7.6 buffer; 8 mM D-glucose; 9.6 mM sodium salt ATP; 19 mM MgCl_2_; 0.1 mM NADP^+^; 5U/ml G6PD enzyme (24). The reaction was initiated by adding 100 μg of extract and monitored for 5 minutes at 28°C. A blank was performed with the same preparation, in the absence of glucose.

Pyruvate Kinase (PK) Activity Assay: Pyruvate kinase activity was measured using a coupled enzymatic assay with lactate dehydrogenase (LDH), by monitoring the decrease in NADH absorbance at 340 nm. In this reaction, PK catalyzes the transfer of a phosphate group from phosphoenolpyruvate (PEP) to ADP, forming ATP and pyruvate. The pyruvate produced is then converted to lactate by LDH, which simultaneously oxidizes NADH to NAD⁺. As NADH is consumed, its absorbance at 340 nm decreases over time, and this decay is used as a readout of PK enzymatic activity. A blank was performed with the same preparation, in the absence of PEP (25). The reaction consists of 39 mM potassium phosphate buffer pH 7.6; 0.58 mM PEP; 0.11 mM NADH; 6.8 mM MgSO_4_·7H_2_O; 1.5 mM ADP; 10 U of LDH; and 250 μg of parasite extract to be measured for activity, in a final volume of 200 μl. The activity was monitored for 5 minutes at 28°C.

Citrate Synthase (CS) Activity Assay: The activity of citrate synthase (CS) was measured based on the decay of Acetyl-CoA in the reaction. Oxaloacetate (OA) was used by CS as a substrate. CS transfers the CoA group to oxaloacetate, generating citrate and Coenzyme A without the acyl group (CoA-SH). A blank was performed with the same preparation, in the absence of OA (26). The reaction consists of 100 μM DTNB; 7.5 mM Tris-HCl pH 8.1; 0.03 mM Acetyl-CoA; 500 μM oxaloacetate; and 100 μg of parasite extract to be measured for activity, in a final volume of 200 μl. The activity was monitored for 5 minutes at 28°C.

Glycerol Kinase (GK) Activity Assay: Glycerol kinase catalyzes the transfer of a phosphate group from ATP to glycerol, resulting in the formation of glycerol-3-phosphate (G3P). The activity of this enzyme was measured at 340 nm from a coupled reaction, adding PK and LDH enzymes to the reaction. G3P has its phosphate group transferred to ADP by the action of PEP, generating ATP and pyruvate. Pyruvate was then converted by LDH into lactic acid, via oxidation of NADH, which is consumed from the medium, resulting in a decreasing absorbance. The reaction consists of 39 mM potassium phosphate buffer pH 7.6; 0.58 mM PEP; 0.11 mM NADH; 6.8 mM MgSO_4_·7H_2_O; 1.5 mM ADP; 10 U of PK/LDH; and 100 μg of parasite extract to be measured for activity, in a final volume of 200 μl. The activity was monitored for 5 minutes at 28°C (27).

Pyrroline-5-carboxylate dehydrogenase (P5CDH) Activity Assay. The activity of P5CDH was measured based on the increase in NADH formation, catalyzed by pyrroline-5-carboxylate (P5C) dehydrogenase, which irreversibly converts NAD^+^-dependent P5C into glutamate. The formation of NADH released by the enzyme is measured at 340 nm (27). The reaction consists of 100 mM KH_2_PO_4_ buffer pH 7.2; 40 mM NAD^+^; 100 μg parasite extract, in a final volume of 200 μl. The activity was monitored for 5 minutes at 28°C. For all enzymatic assays, a control group was included and consisted of reaction solutions without the enzyme substrates to rule out residual activity.

## 3. Results

### 3.1 Cell Cycle Synchronization and Nuclei Isolation: Validation and Overview of Nuclear Proteins Across the Cell Cycle

To decipher the nuclear cell cycle proteins signature, we proceeded with a high resolution labelled quantitative proteomics experiment, detailed in the flowchart at Supplementary Fig1. To synchronize the cell cycle in epimastigotes forms, we used HU that inhibits the ribonucleotide reductase enzyme, blocking the conversion of ribonucleotides into deoxyribonucleotides (dNTPs) leading to a depletion of dNTP pools, effectively preventing DNA synthesis (19). As a result, cells treated with HU are unable to progress through the S phase and are instead halted at the G1/S transition. Thus, cells in S and G2/M were collected upon HU release and cell cycle phase were further confirmed by flow cytometry analysis (Fig 1A).

**Fig 1.**
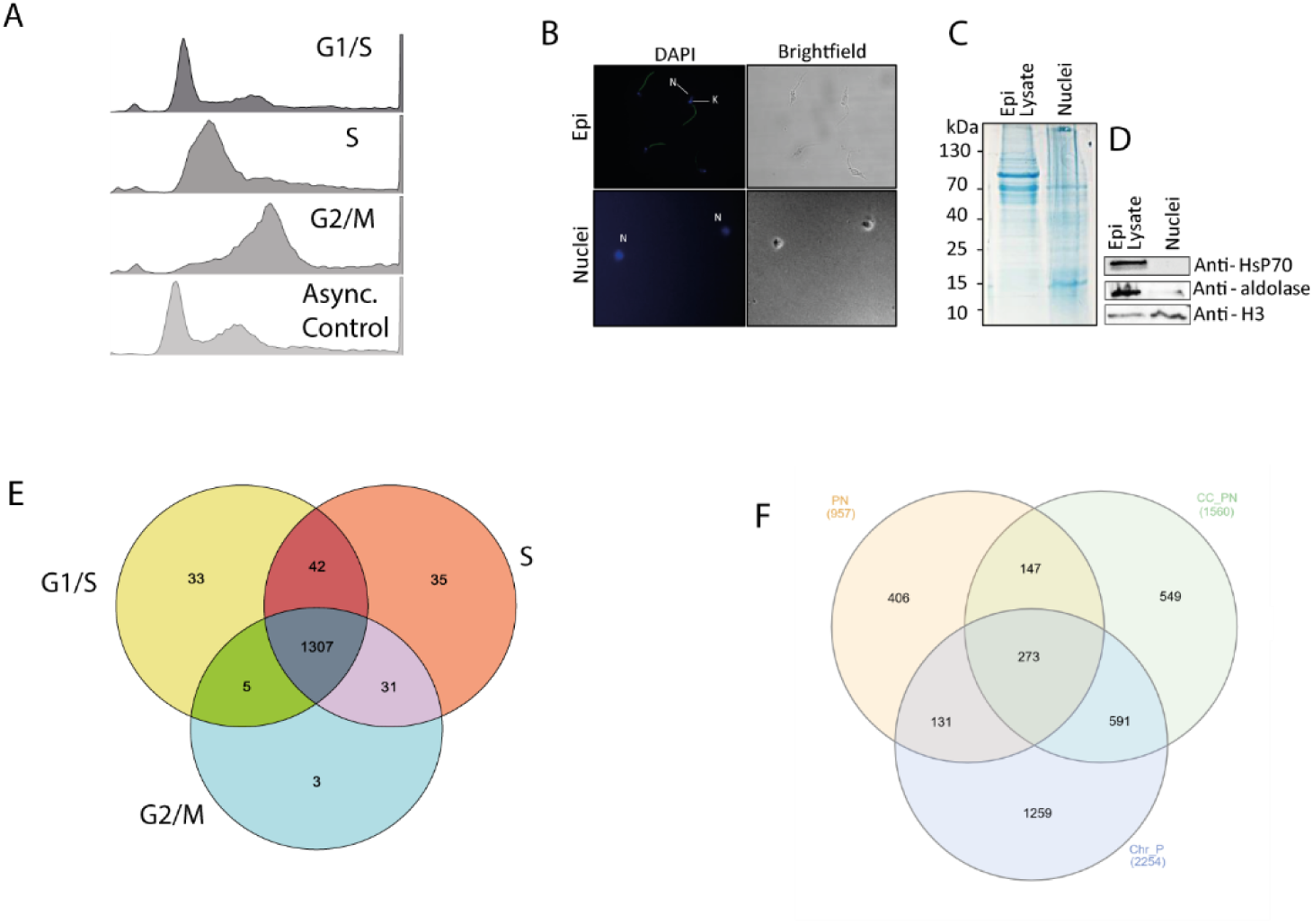
Characterization of Nuclear Protein Extracts and Histone Abundance During the *T. cruzi* Cell Cycle. (A) Representative flow cytometry histograms indicate the distribution of the DNA content in the experimental groups G1/S, S, G2/M and asynchronous control. (B) The nuclear extract fractions were stained with DAPI and visualized under a phase contrast microscope (x100). (C) SDS-PAGE (15%) stained with Coomassie blue, highlighting the enrichment of histones with a molecular weight of ∼15 KDa and (D) representative western blot showing the presence of nuclear marker protein (anti-H3) and the absence of ant-HSP70 (endoplasmatic reticulum), anti-aldolase (glycosomal) in the nuclear fraction and whole cell lysate. (E) Venn diagram of proteins amount throughout the cell cycle. Abundance of canonical (F) and variants (G) histones

Upon cell cycle synchronization, we purified the nuclei and evaluated them by fluorescence microscopy demonstrating the presence of intact nuclear structures without kinetoplast contamination (Fig 1B). The nuclear protein extracts were analyzed by Coomassie-stained gel electrophoresis (Fig 1C) and western blot (Fig 1D). We found that the nuclear extracts were enriched in histone H3 and minimally contaminated with endoplasmic reticulum (HSP70) and glycosome (aldolase) proteins (Fig 1D). As a whole, the results indicated a successful *T. cruzi* nuclear enrichment.

We quantitatively evaluated TMT-labelled nuclear peptides from cell cycle phases (G1/S, S, and G2/M) by LC-MS/MS. We identified 2,937 proteins belonging to 1,560 protein groups (S1 Table). Cellular component analysis from gene ontology (GO) confirmed enrichment in “chromatin” and “chromosome” terms for proteins from all cell cycle phases/transition (S2C Fig), reinforcing the success of our nuclei purification. Principal component analysis (PCA) revealed a distinct separation (explaining 62.3% of the variability) among G1/S, S, and G2/M in the first two components, indicating differences in the nuclear proteomes across cell cycle phases (S2A Fig).

We identified similar numbers of proteins in each cell cycle phase, with more exclusive proteins in S phase (35 proteins), followed by G1/S (33 proteins) and G2/M (3 proteins) (Fig1E) (S2 table). Among those exclusive proteins of G1/S we found signaling and regulatory proteins such as Mitogen-Activated Protein Kinase 10 (MAPK 10) (TcCLB.506229.10) that can play important roles during cell cycle progression (28). Previous nuclear proteomics analysis (De Castro Moreira Dos Santos et al., 2015) identified 957 nuclear proteins, in which 420 (44%) were found on both analyses. Comparison of previous chromatin proteomics retrieved 864 proteins (55%) (Fig 1F).

### 3.4 Histones, nucleotide/nucleoside metabolic proteins and kinases profiling during cell cycle

It is well established that canonical histones, including those from *T. cruzi*, are synthesized in a cell cycle dependent manner, reaching maximal levels in S-phase (29). In contrast, the timing of synthesis for *T. cruzi* histone variants remains unclear, as their expression may not be tightly coupled to DNA replication. As expected, we observed a peak of both canonical and variant protein levels during S-phase (Fig 1F and 1G). On average, their abundance increases more than threefold compared to G1/S phase, followed by a return to near G1/S levels in G2/M for canonical histones and H2B.V. This pattern reflects the expected upregulation of histone synthesis during S-phase, when histones become proportionally more abundant when compared to the total protein content, and their proportional levels decline post-replication as the cell progresses into G2/M. Interesting, the levels of variants histones H2A.Z and H3.V remained relatively high in G2/M suggesting a differential regulation of histone variant dynamics, potentially playing a distinct role in chromatin organization or maintenance after DNA replication. In accordance, we detected two nucleosome assembly proteins (TcCLB.507031.29 and TcCLB.505983.20) that also were increased threefold in S-phase suggesting their involvement in histone deposition.

Nucleotide synthesis is also finely tuned to the cell cycle, ensuring the presence of substrates for DNA synthesis. A multiomics analysis performed in yeast, revealed that nucleotides synthesis peaked in G1 and S phase and proteins associated with nucleobase metabolism are periodically expressed along the cell cycle (30). We detected 30 proteins associated with nucleotide metabolism in our nuclear proteome dataset. Among them, we found enrichment - ranging from 5.7 to 37 times, relative to G2/M-of four proteins in G1/S phase (Fig 2D and 2E), including the nucleoside diphosphate kinase 1 (NDK), the nucleoside phosphorylase, the ribonucleoside-diphosphate reductase small chain (RNR) and the ribose 5-phosphate isomerase (RPI). All these enzymes were related to the production of more substrates (nucleotides and their parts) for DNA synthesis. For example, the enrichment of NDK may increase the conversion of diphosphate nucleotides into triphosphates. RNR catalyzes the conversion of ribonucleotides into deoxyribonucleotides, therefore favoring DNA replication. RPI catalyzes the utilization of ribose-5-phosphate for nucleotide synthesis. Additionally, nucleoside phosphorylase plays a key role in the salvage pathways of nucleosides, which are essential in trypanosomes since they lack the de novo purine synthesis pathway and rely entirely on salvage mechanisms to obtain purines (31,32).

**Fig 2.**
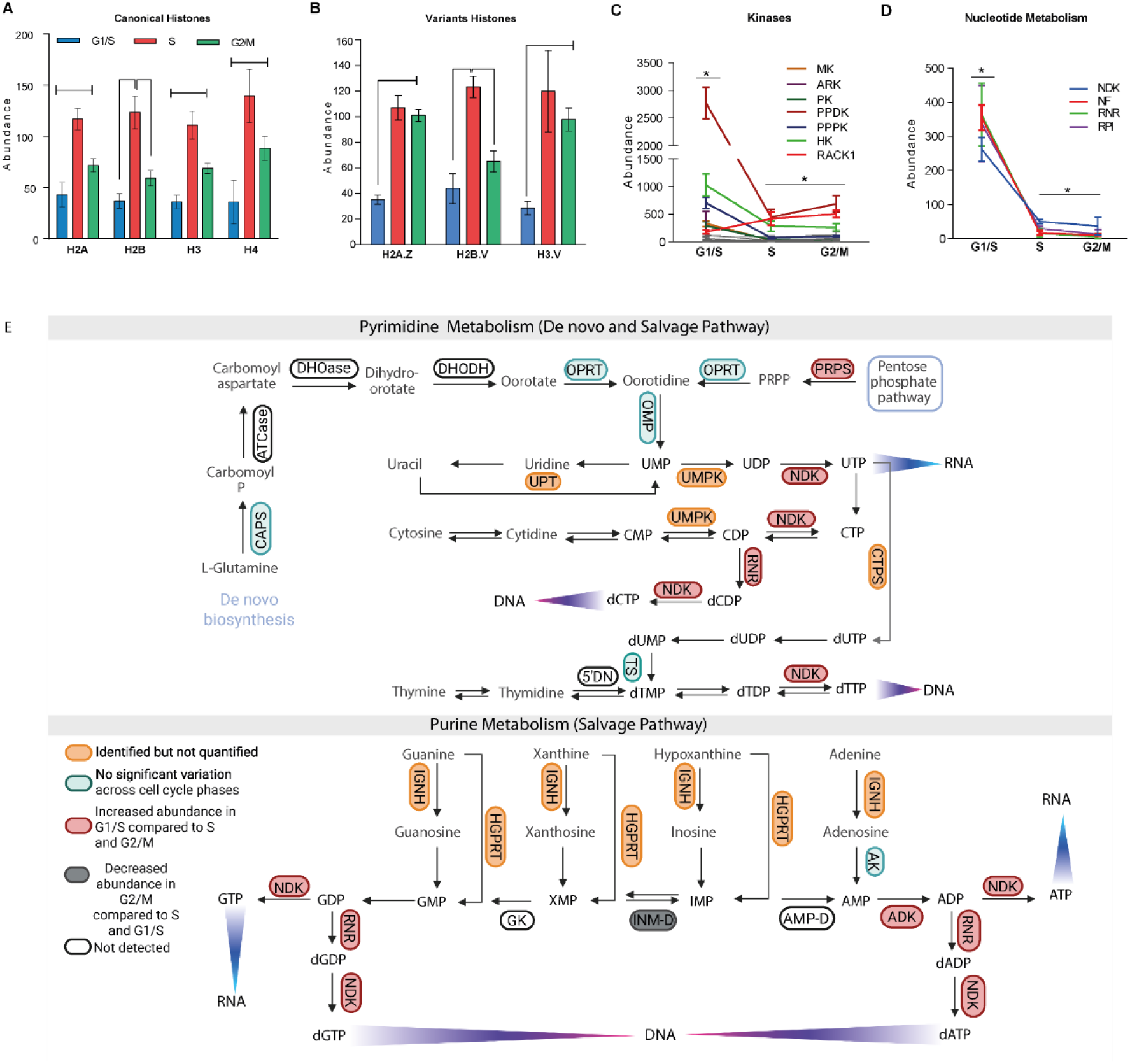
Variation in the Abundance of nucleotide metabolic enzymes and kinases across the *T. cruzi* cell cycle phases. (A) Relative abundance of enzymes associated with nucleotide metabolism (A) and kinases (B) during G1/S, S, and G2/M. Statistical significance was assessed by ANOVA (*p < 0.05, **p < 0.01). (C) Schematic representation of nucleotide metabolism in *T. cruzi*, including both the Pyrimidine *de novo* and Salvage pathways and the Purine Salvage Pathway. Enzymes are color-coded based on their variation throughout the cell cycle: green indicates no significant variation, red denotes increased expression in G1/S, gray represents decreased expression in G2/M, orange corresponds to identified but not quantified enzymes, and black marks enzymes that were not detected. Pyrimidine Metabolism: carbamoyl-phosphate synthetase II (CPSII, EC:6.3.5.5), aspartate carbamoyltransferase (ATCase, EC:2.1.3.2), dihydroorotase (DHOase, EC:3.5.2.3), dihydroorotate dehydrogenase (DHODH, EC:1.3.98.1), orotate phosphoribosyltransferase (OPRT, EC:2.4.2.10), Orotidine-5-phosphate decarboxylase (OMP, EC:4.1.1.23), uracil phosphoribosyltransferase (UPT, EC:2.4.2.9), UMP-CMP kinase 2 (UMPK, EC:2.7.4.22), cytidine triphosphate synthase (CTPS, EC:6.3.4.2), and uridine-cytidine kinase (UCK, EC:2.7.1.48), nucleoside diphosphate kinase (NDK, EC:2.7.4.6), ribonucleoside-diphosphate reductase (RNR, EC:1.17.4.1), 5’-deoxynucleotidase (5’DN, EC:3.1.3.89), thymidylate kinase (TK, EC:2.7.4.9), phosphoribosylpyrophosphate synthetase (PRPS, EC:2.7.6.1). Purine Metabolism: inosine-adenosine-guanosine-nucleoside hydrolase (IGNH, EC:3.2.2.1), hypoxanthine-guanine phosphoribosyltransferase (HGPRT, EC:2.4.2.8), inosine-5’-monophosphate dehydrogenase (INMD, EC:1.1.1.205), guanylate kinase (GK, EC:2.7.4.8), adenosine kinase (AK, EC:2.7.1.20) and adenylate kinase (ADK, EC:2.7.4.3). Additionally, PRPP: phosphoribosyl pyrophosphate, AMP (adenosine monophosphate), ADP (adenosine diphosphate), ATP (adenosine triphosphate), dADP (deoxyadenosine diphosphate), dATP (deoxyadenosine triphosphate), IMP (inosine monophosphate), GMP (guanosine monophosphate), GDP (guanosine diphosphate), GTP (guanosine triphosphate), dGDP (deoxyguanosine diphosphate), dGTP (deoxyguanosine triphosphate), XMP (xanthosine monophosphate), UMP (uridine monophosphate), UDP (uridine diphosphate), UTP (uridine triphosphate), CMP (cytidine monophosphate), CDP (cytidine diphosphate), CTP (cytidine triphosphate), dCDP (deoxycytidine diphosphate), dCTP (deoxycytidine triphosphate), dTDP (deoxythymidine diphosphate), and dTTP (deoxythymidine triphosphate).

In addition to the regulation of nucleotide synthesis, proper coordination of cell cycle events also depends on signaling pathways. Kinases play a pivotal role in the regulation of the cell cycle, orchestrating key events such as DNA replication, mitosis, and cell division (33). We found 42 protein kinases, which were further classified into “Serine/Threonine kinases”, “Glycolysis/Gluconeogenesis Kinases” and “Phosphorylation/Signaling Kinases” (S3 table). Among the kinases, 24% were up-regulated in G1/S (p-value <0.05 and FC>1.5) including hexokinase (TcCLB.510121.20), pyruvate phosphate dikinase (TcCLB.510101.140 and TcCLB.506297.190); Phosphoenolpyruvate carboxykinase (TcCLB.507547.90), nucleoside diphosphate kinase 1 (TcCLB.508707.200), arginine kinase (TcCLB.507241.30), phosphoglycerate kinase (TcCLB.505999.90), homoserine kinase (TcCLB.503597.10) and mevalonate kinase (TcCLB.436521.9)(Fig2C). Notably, we detected upregulation in G2/M vs. G1/S for the receptor for activated C kinase 1 (TcCLB.511211.130), a protein involved in cytokinesis regulation (34), reinforcing the good quality of our results (Fig 2C).

### 3.3 Differentially expressed proteins across the cell cycle

To get further insights into nuclear protein dynamics, we identified 359 proteins exhibiting upregulation and/or downregulation (FDR<0.05, fold changes ≥ 1.5 or ≤ −1.5 and p-value < 0.05) (Fig 3A-C) (S4 table). Comparison between protein expression from G1/S vs. S and G2/M vs. G1/S, retrieved higher number of differentially expressed proteins (DEPs), related to S vs. G2/M. Specifically, in the G1/S vs. S, we detected 100 up-regulated and 78 down-regulated proteins (Fig 3A). Regarding the G2/M vs. G1/S, 84 proteins were down-regulated, while 89 were up-regulated (Fig 3C). Conversely, in the S vs. G2/M, only 8 proteins were differentially expressed (Fig 3B).

**Fig 3.**
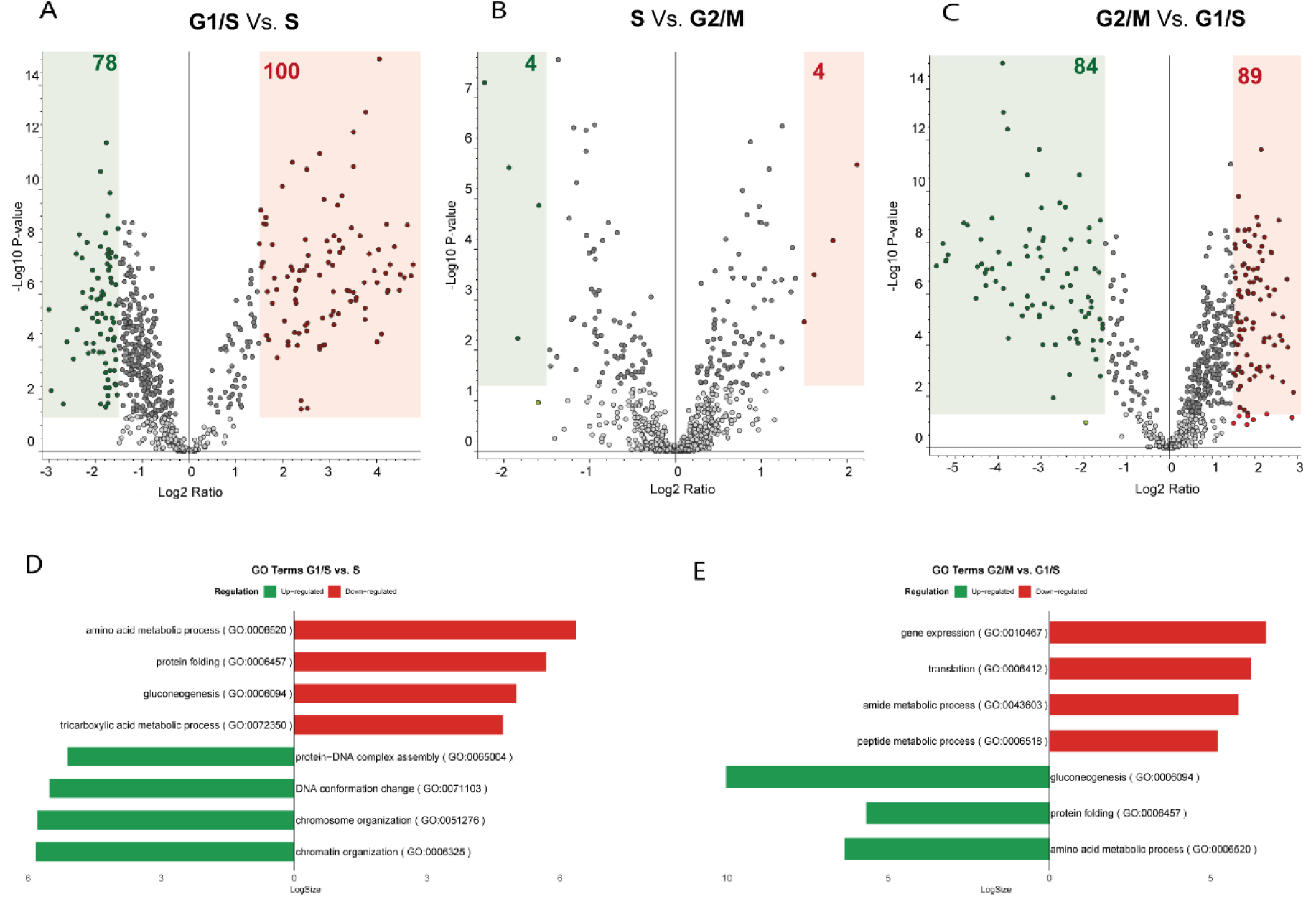
Differentially expressed proteins across the cell cycle. Volcano plots show the differentially abundant nuclear proteins between (A) G1/S vs. S, (B) S vs. G2/M, and (C) G2/M vs. G1/S phases. Proteins with significantly increased (red) (pValue <0.05 and FC>1.5) or decreased (green) (pValue <0.05 and FC<-1.5) abundance are highlighted, and the total number of upregulated and downregulated proteins in each comparison is indicated. (D–E) Bar plots display enriched Gene Ontology (GO) terms from the Biological Process category for the comparisons G1/S vs. S (D) and G2/M vs. G1/S (E). GO terms were obtained from TriTrypDB tools and analyzed using ReviGO, filtering by *p*-value < 0.05. Bar lengths represent the log-scaled size of the GO term groups. Upregulated processes are shown in red, while downregulated processes are shown in green.

To further investigate the biological roles of DEPs across the *T. cruzi* cell cycle, we analyzed enriched GO terms within the “biological process” category (S5 table). The top enriched terms are presented in Fig 3D (G1/S vs. S) and Fig 3E (G2/M vs. G1/S). GO enrichment analysis comparing G1/S and S phases revealed that proteins upregulated in G1 were associated with metabolic and biosynthetic processes, including “amino acid metabolic process” (GO:0006520), “protein folding” (GO:0006457), “gluconeogenesis” (GO:0006094), and “tricarboxylic acid metabolic process” (GO:0072350) (Fig3D). These terms suggest that *T. cruzi* cells are metabolically active during G1, accumulating energy and building blocks required for the upcoming DNA replication. In contrast, proteins more abundant in the S phase were enriched for “chromosome organization” (GO:0051276), “DNA conformation change” (GO:0071103), “protein–DNA complex assembly” (GO:0065004), and “chromatin organization” (GO:0006325) (Fig 3D), indicating that chromatin remodeling and nucleoprotein assembly processes are prominent during active DNA synthesis, as expected.

In the G2/M vs. G1/S comparison, a similar set of metabolic terms remained downregulated in G2/M - “amino acid metabolic process” (GO:0006520), “gluconeogenesis” (GO:0006094), and “protein folding” (GO:0006457)- and thus more enriched in G1/S (Fig3E). Conversely, upregulated proteins in G2/M were associated with “gene expression” (GO:0010467), “translation” (GO:0006412), “amide metabolic process” (GO:0043603), and “peptide metabolic process” (GO:0006518) (Fig3E), suggesting a shift toward increased protein synthesis and gene expression as the cell prepares for mitosis. Together, these results indicate a functional transition across the cell cycle, from a metabolically active G1/S phase to a G2/M phase characterized by upregulation of gene expression and translation-related processes.

### 3.4 Metabolic Compartmentalization

The recurrent enrichment of GO terms related to amino acid metabolism, gluconeogenesis, and the TCA cycle, suggests the presence of metabolic enzymes in the nuclear proteome. This prompted further investigation into their nuclear localization and the potential role of metabolic compartmentalization within the nucleus. The presence of metabolic enzymes within the nuclear compartment was already detected in other organisms (35–37) offering insights about their role within the nuclear environment.

We found that 5.96% (90 proteins) of nuclear proteins were constituted of metabolic enzymes (Fig 4A and S6 Table) enriched with “Fatty acid elongation”, “Citrate cycle (TCA cycle)”, “Pyruvate metabolism” and “Pyrimidine metabolism” according KEGG pathways (p-value < 0.05, Bonferroni) (Fig 4B). To confirm their nuclear location, we compared this list with other *T. cruzi* proteomic datasets related to chromatin (CrP_P) (21) and nuclei (NP_P) (18)(Fig 4C). Interestingly, 28 metabolic proteins were also identified in the studies mentioned above (Fig 5C), including hexokinase (TcCLB.510121.20 or TcCLB.508951.20), pyruvate dehydrogenase E1 component alpha subunit (TcCLB.507831.70), pyruvate dehydrogenase E1 beta subunit (TcCLB.510091.80) pyruvate phosphate dikinase (TcCLB.510101.140), citrate synthase (TcCLB.509801.30), 3- phosphoglycerate kinase (TcCLB.511419.50), which are key enzymes in central carbon metabolism. Forty-four proteins were found both in our current dataset and at chromatin proteomics (Fig 5C). To further confirm their nuclear location, we evaluated their subcellular location using the TrypTag platform (38). We found 23.3% (18/77) of the metabolic proteins identified in our analyses that contains an ortholog in *Trypanosoma brucei* (77 genes) indeed have nuclear localization when the mNeon green tagged parasites were evaluated (S6 table). Among them, proteins associated with the purine and pyrimidine metabolism, inositol phosphosphingolipids phospholipase C (TcCLB.509777.130) and both the alpha and beta subunit of the pyruvate dehydrogenase.

**Fig 4.**
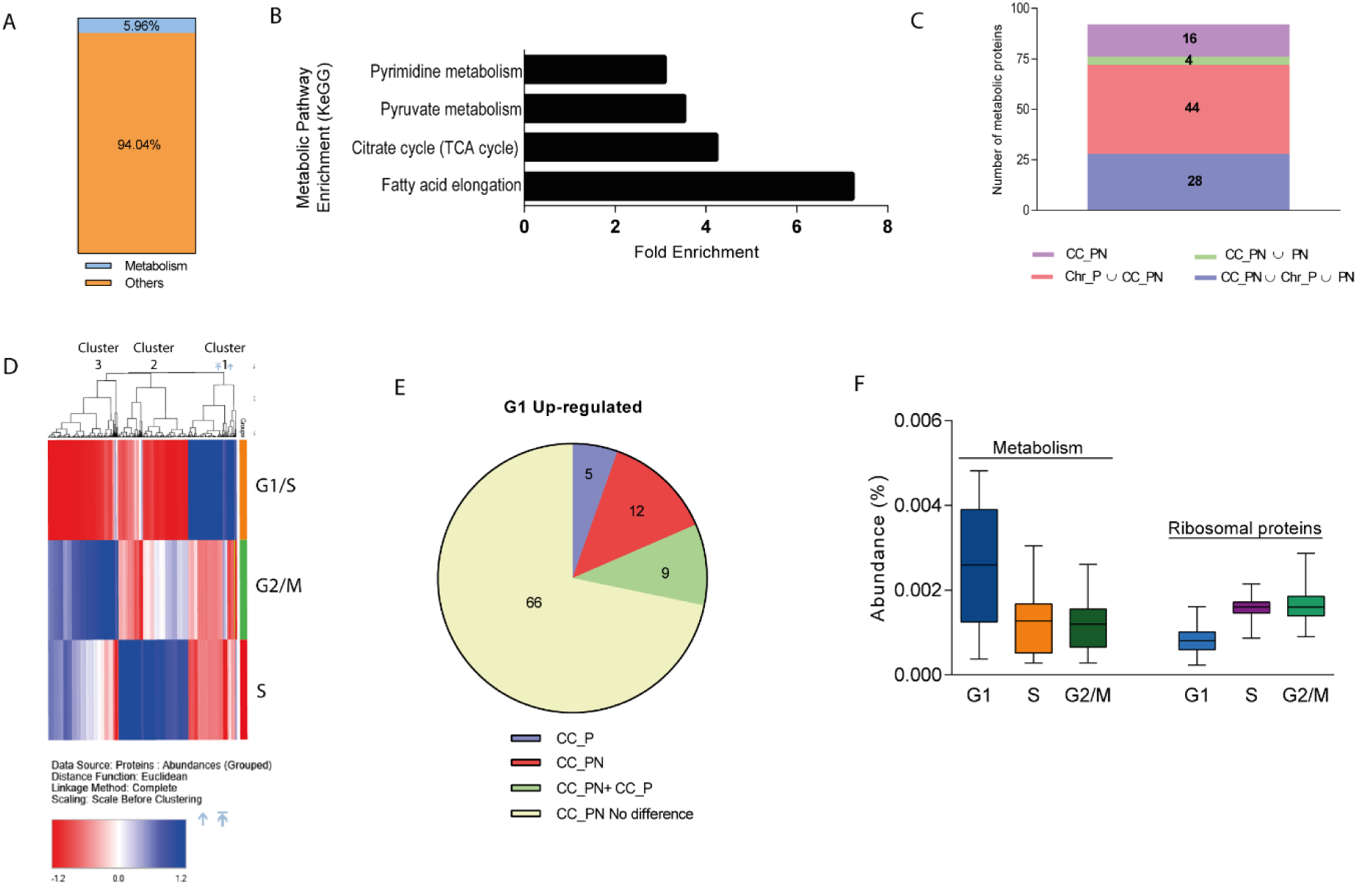
Analysis of nuclear metabolic enzymes across the *T. cruzi* cell cycle. Proportion of nuclear proteins associated with metabolism (5.96%) identified using GO terms related to key metabolic processes, including “cellular amino acid metabolic process” (GO:0006520), “carbohydrate metabolic process” (GO:0005975), “pyruvate metabolic process” (GO:0006090), “ATP metabolic process” (GO:00046034), “nucleotide metabolic process” (GO:0009116), “nucleoside metabolic process” (GO:0009117), “lipid metabolic process” (GO:0006629), and “generation of precursor metabolites and energy” (GO:0006091). (B) KEGG pathway enrichment analysis of the metabolic enzymes identified in (A), performed using TriTrypDB’s tool with a Bonferroni-corrected p-value cutoff of 0.05. Enriched pathways are shown according to fold enrichment. (C) Venn diagram comparing the nuclear proteins identified in this study (CC_PN) with those present in the nuclear proteome (CC_PT) (De Castro Moreira Dos Santos et al., 2015) and those at chromatin proteome (Chr_P) (De Jesus et al., 2017). (D) Hierarchical clustering analysis of all protein expression profiles across different cell cycle phases. The protein data abundance was normalized before clustering by the Euclidean distance ratio. (E) Venn diagram between the nuclear cell cycle proteins identified in this study (CC_NP) and the Cell cycle proteome (CC_P) (Santos Júnior et al., 2021). (F) Box plots representing the relative abundance (%) of nuclear metabolic proteins found in (A) and ribosomal proteins across cell cycle phases.

**Fig 5.**
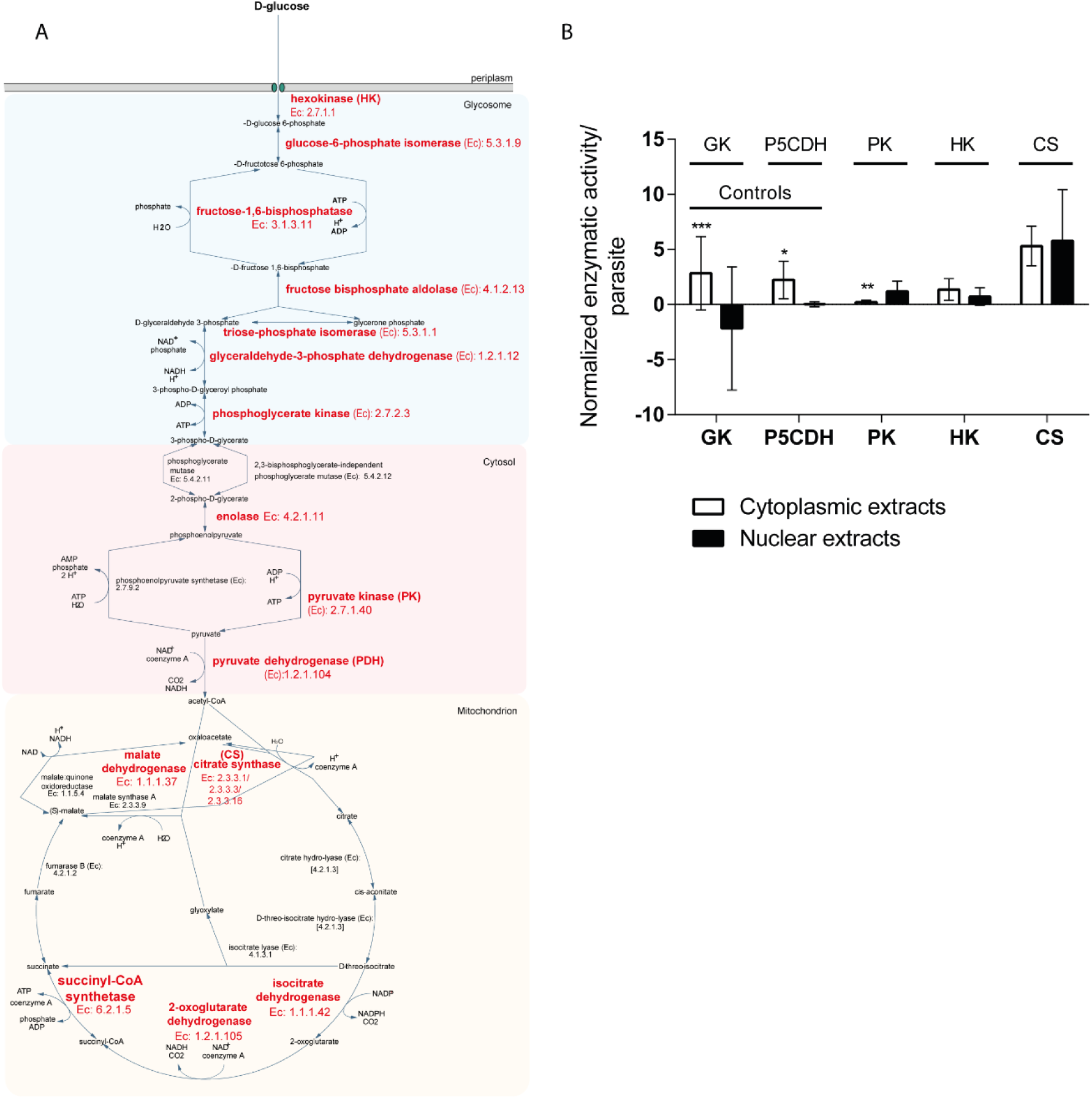
Key enzymes of glycolysis and the TCA cycle exhibit nuclear activity. (A)Activity of hexokinase (HK), citrate synthase (CS), and pyruvate kinase (PK), glycerol kinase (GK) and pyrroline-5-carboxylate dehydrogenase (P5CDH) were obtained using either nuclear and cytoplasmic extracts. The controls used here were not found in chromatin and nuclear proteomes. Reactions without the enzyme substrates were used to rule out residual activity. Enzymatic activities were normalized with the total activities of parasite extract for each listed enzyme. Means ± SD of triplicate data (n=3) are graphically represented. *p < 0.01; **p < 0.001; *** p < 0.0001. One-way ANOVA (Bonferroni’s Multiple Comparison Test). (B) KEGG map of the glycolysis and TCA cycle pathways. Enzymes shown in red were identified in the nuclear proteome throughout the cell cycle.

Molecular traffic between the nucleus and cytoplasm is mediated by nuclear pore complexes (NPCs) in the nuclear envelope. These complexes allow the passage of ions, metabolites, and small proteins (40-60 kDa), but restrict larger macromolecules unless they are bound by nuclear transport receptors, such as importins and exportins (39). Notably, ∼68% (1052) of the nuclear proteins identified in our proteomic analysis were under 60 kDa (S3 Fig), including 74 (82.2%) metabolic proteins, suggesting that their nuclear localization may be facilitated by passive diffusion through NPCs rather than active transport mechanisms.

Temporal compartmentalization of some enzymes can influence cellular function mainly in the dynamic control of cell cycle progression (35). Then, we investigated whether different cell cycle phase/transition were associated with a distinct profile of metabolic protein expression. We found a hierarchical cluster of highly abundant proteins at G1/S compared with S and G2/M that are enriched in metabolic proteins (Fig 4D and S7 Table). To assess whether these metabolic proteins were specifically localized inside the nucleus during the G1 phase or simply upregulated in G1, we compared our dataset with the *T. cruzi* cell cycle proteomics analysis published by Santos Júnior et al. (2021) (Fig 4E). Among the metabolic-associated proteins identified in our nuclear proteome, twelve were only enriched at G1 in our current dataset, including four proteins from the carbohydrate metabolism. Together, these results suggest a specific enrichment of nuclear metabolic proteins specifically associated with the G1 phase, suggesting a possible increase in the nuclear import or retention of metabolic proteins during G1.

To assess whether the abundance of metabolic proteins in nuclear extracts is comparable to that of nuclear proteins, we compared their levels to ribosomal proteins, which are typically abundant in the nucleus. Metabolic enzymes abundance is confirmed to be higher at G1/S phase and lower in S and G2/M phase in contrast with ribosomal proteins that display a progressive enrichment across the cell cycle. Interestingly, metabolic enzymes showed nuclear abundance values in the same order of magnitude as ribosomal proteins across all cell cycle phases (Fig 4F). This comparable abundance suggests that the nuclear presence of metabolic enzymes is not merely due to contamination or passive diffusion but rather may reflect a regulated and potentially functional compartmentalization. The particularly higher abundance of metabolic enzymes in G1 further supports the idea of dynamic metabolic regulation within the nucleus at specific cell cycle stages.

### 3.5 Validation of nuclear metabolic compartmentalization

Given that nearly 6% of our nuclear proteome is composed of metabolic enzymes— primarily those involved in glycolysis, pyruvate metabolism, and the TCA cycle (Fig 4B)— we mapped all identified nuclear enzymes onto central carbon metabolism pathways (highlighted in red, Fig 5A). Our goal was to determine whether these enzymes are confined to a few specific steps or broadly distributed across the pathways. We found that they are indeed widespread, with enzymes present at multiple points throughout both glycolysis and the TCA cycle.

To determine whether they are enzymatically active inside the nucleus, we evaluated the activity of three key metabolic enzymes—hexokinase, pyruvate dehydrogenase E1 component alpha subunit, and citrate synthase—which play essential roles in cellular energy metabolism. Hexokinase is canonically located at *T. cruzi* glycosomes, while the other two at mitochondrion. We assessed their activity in total, cytoplasmic, and nuclear extracts from asynchronous epimastigotes. Of note, the availability of standardized enzymatic assays for these enzymes was also a key factor for their selection. To confirm the absence of cytoplasmic contamination in the nuclear extracts, we also assessed the nuclear activity of enzymes not identified in nuclear proteomes (current dataset and from De Castro Moreira Dos Santos et al., 2015). Specifically, we analyzed glycerol kinase (GK) and pyrroline-5-carboxylate dehydrogenase (P5CDH), which are primarily localized in glycosomes and mitochondria, respectively.

Our analysis confirmed the presence and the nuclear activity of hexokinase (TcCLB.508951.20 and TcCLB.510121.20), pyruvate kinase (TcCLB.507993.390), and citrate synthase (TcCLB.509801.30) (Fig 5B, supplementary Fig 4). In contrast, P5CDH and GK -both negative controls, as expected, displayed no activity in the nuclear extracts.

These results confirm the presence of key enzymes involved in cellular energy metabolism within the nucleus and further demonstrate their metabolic activity. This suggests that these enzymes are not passive contaminants generated during experimental approaches but may be translocated to the nucleus likely performing specific functions.

## 4. Discussion

Our study provides a comprehensive analysis of nuclear proteome dynamics during the cell cycle of *T. cruzi*. Using synchronized epimastigote forms, we isolated nuclei and extracted their proteins identifying and quantifying more than one thousand proteins. A substantial number of DEPs were identified across cell cycle phases, reflecting the dynamic nature of nuclear processes during cell cycle transitions. Up-regulated proteins in the G1/S phase were mainly associated with energy production and metabolism, while those in the G2/M phase were linked to protein synthesis and cell division. Notably, our study highlighted the presence and activity of metabolic enzymes within the nucleus, with enzymes related to carbohydrate, aminoacid, lipid, nucleotide metabolism and energy production, being upregulated, particularly during the G1/S. These results suggest a link between nuclear metabolic activity and cell cycle regulation.

The primary challenge for nuclear proteome analysis is the isolation of pure nuclei, minimizing cytoplasmic contamination. In this study, we implemented a robust protocol that combined cell membrane lysis, ultracentrifugation, and organelle fractionation to isolate intact nuclear structures (20). The purity of the nuclear fraction was confirmed using multiple approaches, including DAPI staining for direct visualization of nuclear structures under microscopy, histone enrichment analysis by SDS-PAGE, and the absence of glycosomal and endoplasmic reticulum markers in western blot assays. Additionally, gene ontology (GO) enrichment analysis in the “cellular component” category further validated the successful isolation of nuclear proteins. In addition, our data analysis revealed significant overlap of GO terms found in other eukaryotic cell cycle proteomic analysis (40,41) suggesting that essential nuclear processes are maintained through evolutionary constraints. Despite the evidence supporting the successful purification of nuclear extracts, we cannot entirely exclude the possibility of minor non-nuclear contamination.

Our nuclear proteomics analysis confirmed the expected increase of canonical histones during S-phase. Importantly, our data revealed for the first time that the histone variants H2A.Z, H2B.V, and H3.V also show a significant increase in S-phase, suggesting that their expression or regulation may be coupled to DNA replication, similar to canonical histones. In contrast, H3.V and H2A.Z maintain relatively stable levels after S-phase (when normalized to total nuclear protein), indicating a proportional enrichment during G2/M phases — a pattern opposite to that of canonical histones and H2B.V. This suggests that these variants may continue to be expressed and incorporated into chromatin beyond S-phase, potentially playing roles in cell cycle progression. H3.V and H2A.Z are known to be located at transcription termination and initiation sites, respectively (42). Despite forming dimers (e.g., H2A.Z with H2B.V) (42,43) the distinct expression profiles of these variants suggest they may serve different functions throughout the cell cycle or be incorporated into chromatin at distinct genomic loci such as damage DNA, possibly in combination with canonical histones. These observations open new avenues for understanding the specific roles of histone variants in trypanosomatids. Given that our analyses are based on nuclear extracts, we infer that their enrichment during S-phase likely reflects chromatin deposition, although we cannot entirely exclude their presence in other nuclear substructures such as the nuclear matrix.

Advances in mass spectrometry have enabled the identification and quantification of an increasing number of proteins, significantly enhancing the depth of proteomic studies. Leveraging these advances, our study identified a greater number of nuclear proteins compared to the previous nuclear proteomics report (18), and obtained quantitative data by using TMT labeling. These methods combined further improved sensitivity and accuracy in protein identification/quantitation, which is essential for detecting low-abundance nuclear proteins that may play critical roles in cell cycle regulation. Among those low abundant proteins are the cyclins, cyclin-dependent kinases (CDKs), and other kinases. These proteins are crucial regulators of the cell cycle, driving its progression by phosphorylating key substrates involved in cellular growth and division (33,44). Here, we did not detect any cyclins or CDKs in our dataset, although some of them may be located at the nucleus. It is worth noting that they were also absent from previous cell cycle proteome and transcriptome data (16,17) suggesting that they are indeed very low abundant proteins/transcripts.

Nevertheless, we identified a substantial number of kinases. Among the 42 kinases detected, we observed distinct patterns of abundance across the cell cycle phases. Only one kinase, the receptor for activated C kinase 1 (RACK1), was upregulated in G2/M, which aligns with its proposed role in cytokinesis control (34). A considerable proportion of these kinases (∼24%) showed significant upregulation in G1/S, suggesting a potential role in preparing the parasite for DNA replication. Notably, this group included several metabolic kinases, such as hexokinase, pyruvate phosphate dikinase, and phosphoenolpyruvate carboxykinase, which may reflect an increased energy demand and biosynthetic activity required for cell cycle progression. In contrast, a subset of kinases maintained stable abundance levels throughout the cycle, suggesting constitutive roles unrelated to cell cycle regulation. These differential expression patterns highlight a possible phase-specific contribution of kinases to nuclear processes and cell cycle regulation in *T. cruzi*.

Metabolic enzymes, particularly those related to energy metabolism, are typically associated with processes occurring in the cytoplasm, mitochondria, or in case of trypanosomes, also in glycosomes, rather than within the nucleus itself (40,41). However, several enzymes associated with glycolysis and the TCA cycle have already been reported to be localized within nuclear compartments (36,37,45,46). Speculations about their nuclear function include not only the performance of their canonical roles, such as contributing to the regulation of the nuclear NAD⁺/NADH or ATP pool, but also potential non-canonical functions. These may involve influencing the abundance of epigenetic marks on both histones and DNA, acting as transcriptional regulators of specific genes, or participating in DNA repair processes (34). In yeast cells, nuclear-localized metabolic enzymes influence epigenetic regulation (36). For instance, the nuclear pyruvate dehydrogenase complex generates acetyl-CoA for histone acetylation, a key step in epigenetic modification (36,37). Additionally, mitochondrial enzymes that transiently enter the nucleus have been implicated in epigenetic remodeling during zygote activation (47). The identification of metabolic proteins within the *T. cruzi* nucleus, together with their differential abundance across cell cycle phases — reflecting spatial-temporal regulation — provides new insights into how *T. cruzi* coordinates nuclear metabolism to meet the specific demands of its epigenetic machinery. In line with this, we recently observed dynamic changes in histone post-translational modifications (PTMs) throughout the cell cycle (48), further supporting the idea that epigenetic requirements fluctuate during cell cycle progression.

The detection of the E1 subunits (α and β) of the pyruvate dehydrogenase (PDH) complex within the nuclear proteome raises intriguing questions about their potential role beyond canonical mitochondrial metabolism. Their nuclear location was also detected at TrypTag platform. While the E1 component catalyzes the decarboxylation of pyruvate, the complete conversion to acetyl-CoA requires the sequential activity of the E2 (dihydrolipoamide transacetylase) and E3 (dihydrolipoamide dehydrogenase) subunits, which were not detected in our nuclear dataset. This suggests that the classical PDH reaction is unlikely to occur fully within the nucleus. Nevertheless, the presence of E1 could reflect non-canonical functions, such as the generation of metabolic intermediates or moonlighting functions unrelated to catalysis. Further experimental validation will be required to determine whether nuclear E1 forms functional complexes, interacts with alternative partners, or contributes to local acetyl-CoA pools via yet unidentified pathways.

Our findings provide a new perspective on how *T. cruzi* may coordinate nuclear and metabolic activities, suggesting a streamlined adaptation that could be crucial for the parasite’s survival and proliferation during its cell cycle. Further research into the nuclear proteome of *T. cruzi* and its interplay with metabolism and epigenetics will be essential for understanding the unique biology of this parasite and may uncover potential targets for therapeutic intervention. In addition to proteins with known functions, our dataset also revealed several hypothetical proteins within the nuclear fraction. Although their functions remain uncharacterized, the information on their subcellular localization and dynamic changes throughout the cell cycle provides valuable insights that may guide future studies and contribute to a better understanding of their biological roles.

## 5. Conclusion

The study of the nuclear cell cycle proteome through cell cycle synchronization and nuclear isolation in *T. cruzi* has provided valuable insights into the complex dynamics of nuclear proteins. By employing advanced proteomic techniques, we identified a broad spectrum of nuclear proteins, revealing distinct expression patterns across different cell cycle phases-G1/S, S, and G2/M.

Our quantitative analysis demonstrated clear phase-specific variations in protein abundance, with differential expression patterns highlighting significant changes between phases. In particular, the G1/S phase was characterized by increased energy production and metabolic enzyme activity, while proteins involved in DNA packing marked the S phase. The G2/M phase showed elevated levels of proteins associated with protein synthesis and microtubule polymerization, critical for cell division. Additionally, the profiling of nuclear kinases revealed that many kinases are upregulated during specific phases, reflecting their crucial role in cell cycle regulation.

The observation of key metabolic enzymes, particularly in the G1/S phase, present and active in the nucleus emphasizes the importance of metabolic nuclear compartmentalization. These findings highlight a possible involvement of metabolic enzymes and/or their metabolites in the modulation of nuclear processes in *T. cruzi*.

## Author Contributions

A.P.J.M obtained the histone extracts, performed mass spectrometry analysis and processing. C.G.C performed enzymatic activity assays. S.M. acquired the mass spectrometry data. A.P.J.M and J.P.C.C. performed data analysis. A.P.J.M., C.G.C., R.B., A.M.S., M.C.E. and J.P.C.C. contributed to the writing, and rationale of this article. All authors contributed to the revision of this work.

## Acknowledgments

We thank Ivan Novaski Avino, Karin Navarro and Ismael Silva for technical assistance. We thank Dr. Alison Alencar for their valuable discussions on mass spectrometry analysis. We thank the NBBC-HPC/Ibu for computational support. We are grateful to VeuPathDB for providing an excellent platform for Trypanosome data analysis.

## Data availability

The RAW files can be accessed at PRIDE PXD064848.

## Disclaimer

This manuscript was reviewed using ChatGPT to assist with grammar, spelling, and clarity improvements. All scientific content, interpretations, and conclusions remain the sole responsibility of the authors.

## Supporting information

**S1 Fig.**
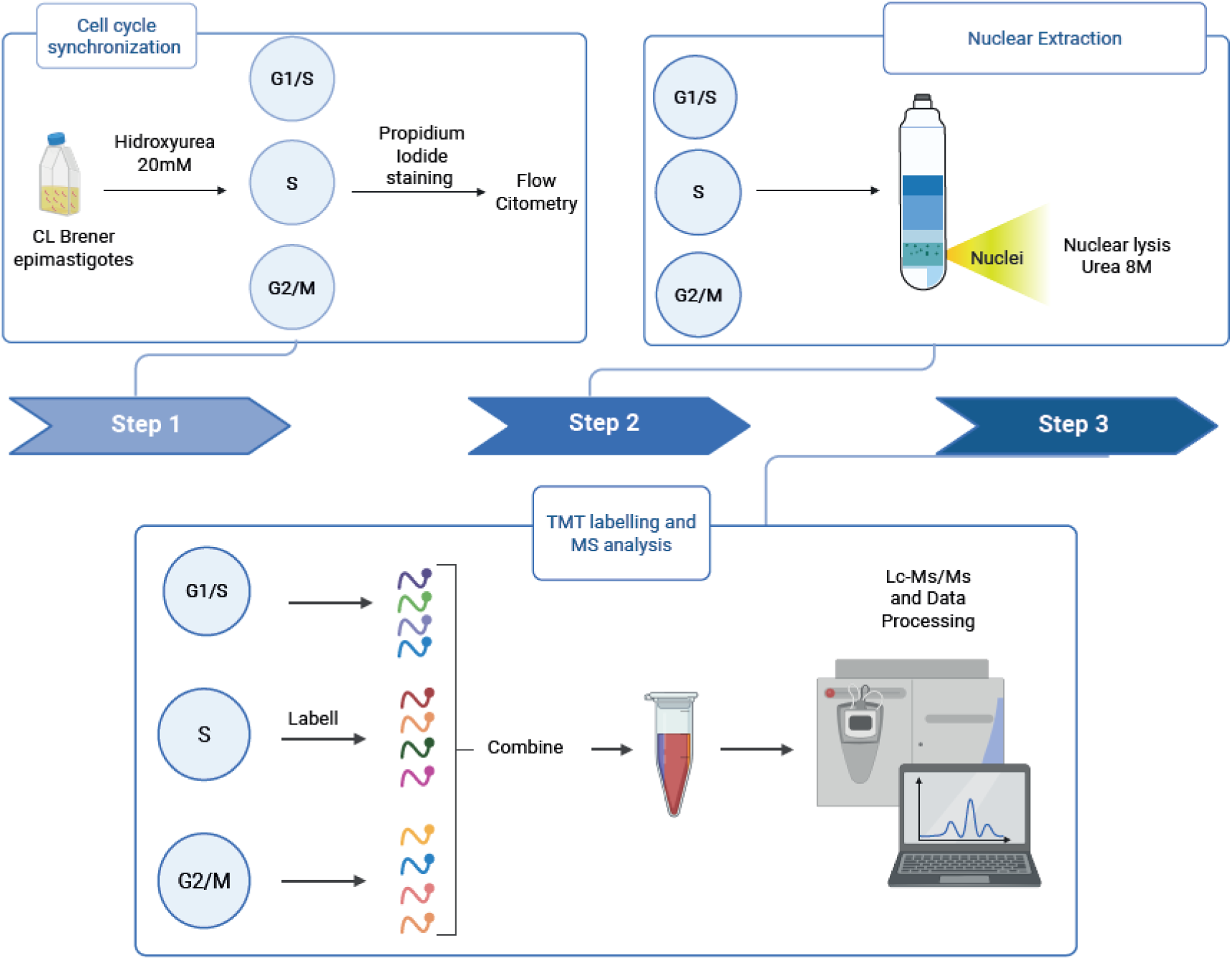
Integrated workflow for cell cycle synchronization and nuclear proteome profiling in T. cruzi. Workflow to cell cycle synchronization and nuclear proteome. Exponential epimastigotes cultures (CL Brener) were synchronized in G1/S, S and G2/M using 20mM of hydroxyurea for 24h(HU). The cell synchronization was confirmed by PI staining and flow cytometry. The nuclei extraction, was realized followed Marques Porto, 2002. Briefly, parasites were lysed parasites with sucrose and NP-40 solutions. After sequentially washing to remove the detergent residues, the pellet was put on percoll gradient. Through ultracentrifugation (65000 x g – 120min), the nucleus was obtained and lysed with an 8M urea solution to release the nuclear proteins. Nuclear protein extracts were cleaned and digested with FASP™ Protein Digestion Kit (Expedeon-Abcam-USA) and labelled with TMT® (Thermo-TMT 16plex Mass Tag Labeling kit cat. 90110) following the manufacturer’s recommendation. Mass spectrometry was performed proceeded using Orbitrap Fusion Tribrid. The TMT data was processed in Thermo Scientific Proteome Discoverer, software version 2.5, with SEQUEST® searc.

**S2 Fig.**
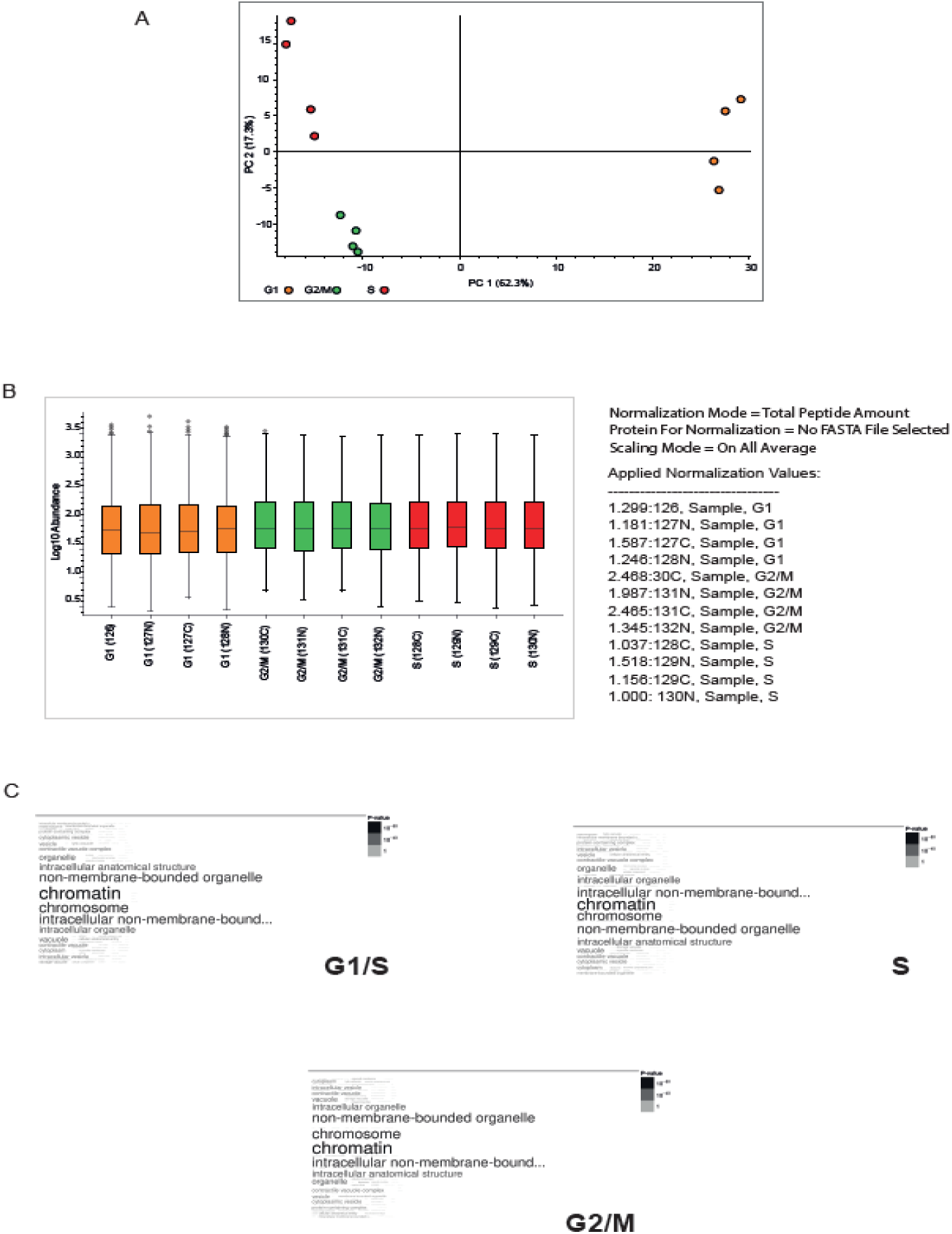
Nuclear proteome dynamics: PCA clustering, protein abundance profiling, and GO-based nuclear enrichment analysis. (A) Principal-component analysis of quadruplicate samples in G1/S (orange), S (red) and G2/M (green). (B) Representation of the normalized relative protein abundance profile obtained through TMT analysis. Each bar corresponds to a replicate and reflects its abundance in the experimental samples. Error bars represent standard deviations from quadruplicate measurements. (C) Enrichment of nuclear proteins by Gene Ontology (GO) analysis. Word cloud demonstrates the enrichment of terms in the cellular components category (GO) for groups G1/S, S, and G2/M, respectively. The nuclei enrichment was evaluated using the TritrypDb, and the significance cutoff for the p-value <0.05. Font sizes in the word cloud are proportional to -log10 of the adjusted p-value for each enriched GO terms.

**S3Fig.**
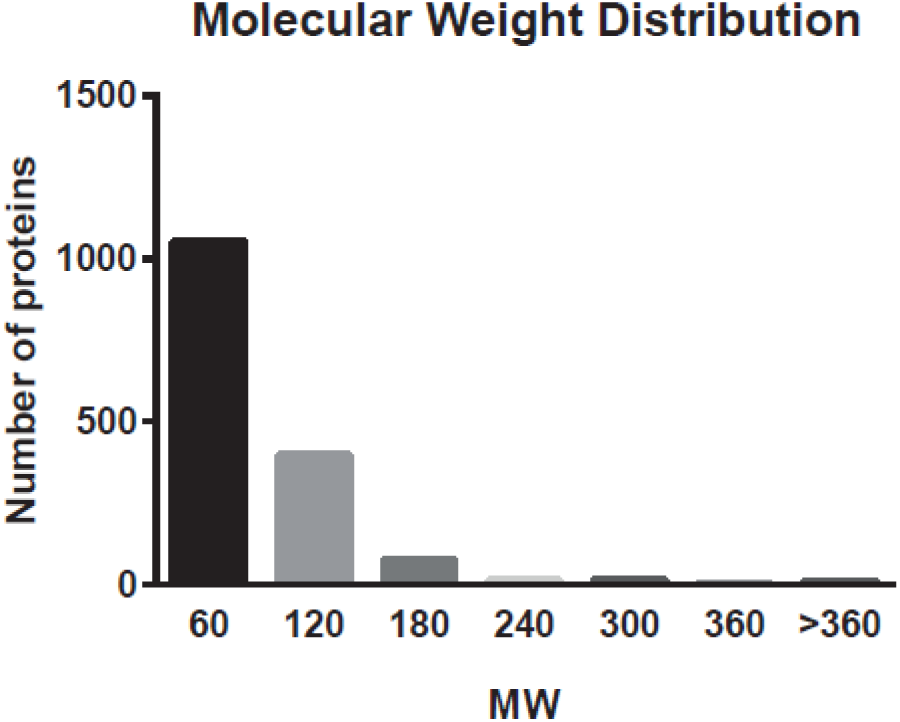
Molecular weight distribution of nuclear proteins (kDa).

**S4 Fig.**
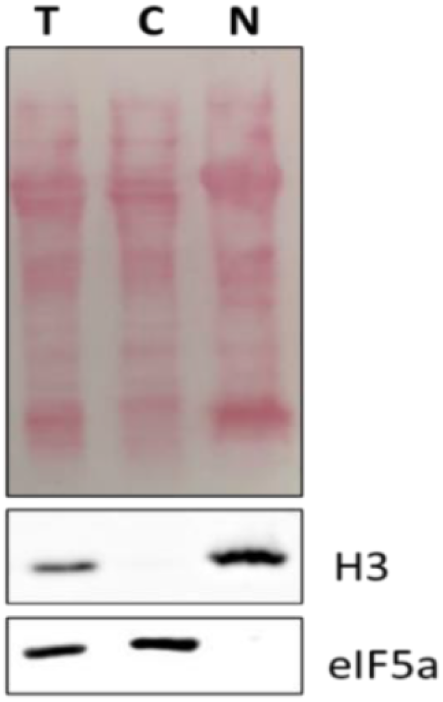
WB showing that nuclear and cytoplasmic separation for enzymatic activity assays. Western blotting of T. cruzi CL Brener WT cells showing labeling with anti-eIF5a (1:5000, mouse), a marker of the cytoplasmic fraction, and anti-histone H3 antibody (1:2000, rabbit), a marker of the nuclear fraction. The protein profile is visualized with Ponceau staining in red. T – Total extract; C – Cytoplasm; N – Nucleus. Representative result of three independent experiments (n=3).

